# Diet-Induced Obesity Exacerbates *Helicobacter pylori*-Associated Precancerous Phenotypes

**DOI:** 10.64898/2026.07.10.737708

**Authors:** Xuyao Zhao, Nicholas Wojcicki, Kee-Hong Kim, Nadia Atallah Lanman, Viju Vijayan Pillai, Valerie Phoebe O’Brien

## Abstract

Stomach infection with the bacterium *Helicobacter pylori* (*Hp*) can cause chronic gastric inflammation, metaplasia (transdifferentiation of mature cell types), dysplasia (abnormal cells), and finally cancer. Obesity can also increase gastric cancer risk. However, host-*Hp* interactions during obesity are poorly understood. Here we investigated the impact of diet-induced obesity in two mouse models of *Hp*-associated disease. To model chronic gastric inflammation, we used C57BL/6 mice, and to model more severe disease, we used transgenic mice in which tamoxifen induces gastric expression of a constitutively active *Kras* allele, leading to metaplasia. We fed mice a high-fat diet (60% kilocalories from fat) to induce obesity, or a matched control diet (10% kilocalories from fat), then infected them with *Hp* or mock-infected them. In mock-infected C57BL/6 mice, high-fat diet had a minimal impact on gastric pathology and gene expression. In *Hp*-infected C57BL/6 mice, high-fat diet increased inflammation at the junction between the glandular stomach and non-glandular forestomach, a squamous epithelium similar to the human esophagus, and increased gastric expression of the cancer-associated genes *Cldn7* and *Reg3g*. In KRAS+ mice with or without *Hp* infection, the impact of diet-induced obesity was more apparent, with increased pseudopyloric metaplasia and dysplasia (abnormal cells). As well, high-fat diet caused an expansion of metaplastic pit cells, a lineage we previously found to be associated with *Hp*-driven inflammation. Thus, in these mouse models, diet-induced obesity does not directly drive gastric immunopathology, but enhances the development of precancerous changes under susceptible conditions.

**IMPORTANCE:** Most gastric cancers are caused by stomach infection with the bacterium *Helicobacter pylori*. However, most infected individuals never develop cancer. Therefore, additional risk factors must tip the balance toward gastric cancer development. Obesity, or excessive body fat accumulation that poses a risk to health, is associated with gastric cancer development. However, specific mechanisms for obesity-driven gastric cancer risk are not well defined. Here we tested the hypothesis that obesity would exacerbate *Helicobacter pylori*-associated disease phenotypes using two clinically relevant mouse models. In wild-type mice, obesity induced by a very high-fat diet had a minimal impact on the stomach in the absence of infection, but increased the expression of some cancer-associated genes during infection. However, in mice with genetically driven precancer, diet-induced obesity exacerbated the disease pathology, especially in infected mice. Therefore, obesity’s impact on gastric cancer risk may be more evident in the later stages of the disease.

## INTRODUCTION

About 80% of gastric cancer cases are attributed to *Helicobacter pylori* (*Hp*) stomach infection ^1, 2^. This bacterium infects half the world and causes gastric inflammation (gastritis), which in some individuals can cause gastric atrophy (loss of appropriate gastric glands), metaplasia (conversion of mature gastric epithelial cells to other cell types), dysplasia (presence of abnormal cells), and finally gastric cancer ^3–9^. Metaplasia in the stomach includes pseudopyloric metaplasia, where native glands of the oxyntic gastric mucosa are replaced by mucus-secreting glands similar to those of the distal stomach, as well as gastric intestinal metaplasia, where native glands are replaced by intestinalized glands ^10^. *Hp* infection elicits a strong immune response that controls but does not eradicate the infection ^11–13^. The bacterium also possesses virulence factors that can directly damage host cells, promoting carcinogenesis ^14^. However, the specific mechanisms through which *Hp* infection drives metaplasia and gastric cancer development are still incompletely understood.

Obesity, defined as excessive body fat accumulation that poses a risk to health, is another major public health challenge that affects over 40% of U.S. adults ^15^. Obesity is associated with numerous gastrointestinal complications ^16^, including gallstones, cirrhosis, gastroesophageal reflux, stomach metaplasia ^17^, and several cancers, including gastric cancer ^16, 18–28^. Obesity’s impact on cancer risk is often attributed to chronic systemic inflammation and systemic metabolic dysregulation that can promote tumor growth ^29^. It is well recognized that enlarged and inflamed adipose tissue in obese individuals contributes to chronic inflammation, both systemically and within the gastrointestinal tract ^30–34^. Obesity also impairs proper immune cell functions ^35^, including reducing the ability of neutrophils to kill bacterial pathogens ^36, 37^. As well, diet-induced obesity impairs how epithelial cells respond to injury ^38^ and delays wound healing ^39^. Finally, high-fat diet-induced obesity can reprogram epithelial cells, promoting stemness and tumorigenic potential ^40, 41^. Despite these observations, the molecular consequences of diet-induced obesity on the stomach are poorly understood. While several epidemiological studies have reported a high prevalence of gastritis in obese subjects ^31, 32^, molecular investigation of the main gastric cancer risk factor, *Hp* infection, in the context of obesity is lacking.

Here we used a high-fat diet (60% kilocalories from fat) to induce obesity in mice, then infected them with *Hp* to assess whether obesity would impact host-pathogen interactions in the stomach. To model the impact of obesity on an otherwise healthy stomach environment, we used C57BL/6 mice. To model the impact of obesity when the stomach harbors preneoplastic changes, we used the *Mist1-Kras* transgenic mouse model of gastric pseudopyloric and intestinal metaplasia. These mice have a tamoxifen-inducible Cre recombinase under the control of the *Mist1* promoter. Subcutaneous tamoxifen induces KRAS^G12D^ in *Mist1*-expressing cells, which in the stomach are progenitors and chief cells ^42, 43^. After six weeks, these “KRAS+ mice” exhibit pseudopyloric metaplasia and mild dysplasia ^42^. We previously found that concomitant *Hp* infection (“*Hp*+KRAS+ mice”) significantly exacerbated disease in this model ^44^, with enhanced inflammation and the expansion of inflammation-associated metaplastic pit cells that express the intestinal mucin *Muc4* ^45^. We found that in C57BL/6 mice, diet-induced obesity did not elicit metaplasia directly. However, in *Hp*-infected mice, high-fat diet increased gastric inflammation at the limiting ridge (the demarcation between the mouse glandular stomach and non-glandular forestomach) and induced expression of the cancer-associated genes *Cldn7* and *Reg3g*. In the transgenic metaplasia model, high-fat diet caused increased disease pathology, including hyperplasia, pseudopyloric metaplasia, and dysplasia, in both KRAS+ and *Hp*+KRAS+ mice, and promoted the expansion of metaplastic pit cells. Together, these results suggest that obesity may contribute to gastric cancer risk by exacerbating *Hp*-driven metaplasia phenotypes.

## RESULTS

*Hp* is a well-established gastric carcinogen, and obesity is a known risk factor that contributes to gastric cancer development. Therefore, we reasoned that host-pathogen interactions might be altered in the stomach during obesity. To explore this idea, we used a high-fat diet (60% kilocalories from fat) to rapidly induce weight gain in C57BL/6 mice (see **Methods** and **Supplementary Methods**). Male and female weanlings were randomly assigned to receive either high-fat diet or a matched standard diet (10% kilocalories from fat), which they remained on for the duration of the experiments. After 15 weeks on the diet to induce obesity, as indicated by body weight (**Fig S1A**), mice were infected with *Hp* or mock-infected. Gastric disease phenotypes were assessed at 12 weeks post-infection (27 weeks on the indicated diet).

### Diet-Induced Obesity Did Not Directly Elicit Gastric Disease

To gain a global view of changes that could occur in the stomach due to high-fat diet and/or *Hp* infection, we assessed gastric pathology in formalin-fixed, hematoxylin and eosin-stained tissue sections (see **Supplementary Methods** for a description of the scoring criteria). We first assessed the effects of high-fat diet on the gastric corpus epithelium (**Fig 1A**) and limiting ridge (**Fig 1B**). Compared to mock-infected mice fed a standard diet, mock-infected mice fed high-fat diet had no gross histopathological changes (**Fig 1A**); the mice all had the expected cellular composition of gastric glands, with pit, parietal, neck, and chief cells all present and no evidence of metaplasia (precancer). As well, there was no overt mucosal inflammation; a blinded analysis of immune cell infiltration revealed a median score of 0 (no inflammation) in mock-infected mice, regardless of diet (**Fig 1C**). In contrast, 12 weeks of *Hp* infection in mice fed a standard diet led to a substantial tissue thickening, suggesting hyperplasia. As well, these mice had increased immune cells in the gastric lamina propria, with a median mucosal inflammation score of 1. However, high-fat diet feeding did not further increase the tissue pathology or the extent of mucosal inflammation in *Hp*-infected mice, with a median mucosal inflammation score of 1 in this group as well.

**Figure 1.**
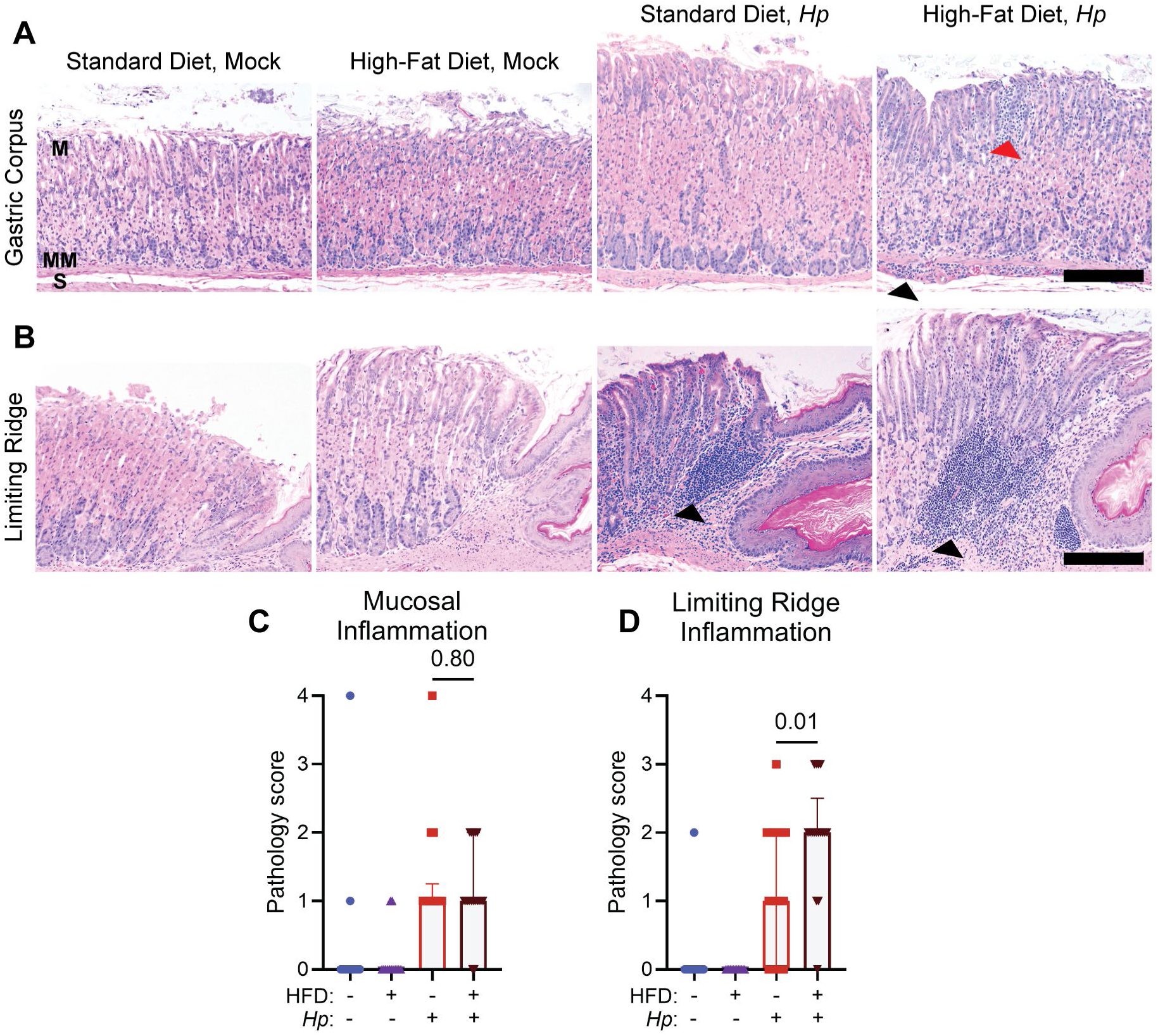
High-fat diet does not induce substantial gastric tissue pathology. C57BL/6 weanlings were randomly assigned to receive high-fat diet (HFD) or standard diet. After 15 weeks to induce obesity, mice were infected with *Hp* or mock-infected for 12 weeks. Mice were then euthanized and a blinded analysis of tissue pathology was performed. **A)** Representative images of the gastric corpus are shown. M, mucosa; MM, muscularis mucosa; S, submucosa. Red arrowhead shows an example of inflammatory cells in the mucosa and black arrowhead shows an example of inflammatory cells in the submucosa. **B)** Representative images of the limiting ridge (junction between glandular stomach and non-glandular forestomach) are shown. Black arrowheads show examples of mononuclear inflammatory cell aggregates. **C)** Inflammation in the gastric corpus mucosa was quantified. **D)** Inflammation at the limiting ridge was quantified. Scale bars, 100 µm. Statistical significance was assessed using the Mann-Whitney U test. Dots represent individual values for each mouse, bars indicate the median and error bars indicate the interquartile range. Data are from n=13-18 mice per group assessed in N=2 independent mouse experiments.

Obesity is strongly associated with cardia cancer, which occurs at the top of the stomach near the esophagus ^18, 19^. Therefore, we specifically assessed inflammation at the limiting ridge, the demarcation between the non-glandular forestomach and glandular stomach (akin to the Z-line in the human stomach at the gastroesophageal junction). In mock-infected mice fed a standard diet or high-fat diet, the appearance of the limiting ridge was unremarkable (**Fig 1B**). *Hp* infection caused increased inflammation at the limiting ridge, with a distinct lymphoid aggregate present in 7 of 17 mice and a median limiting ridge inflammation score of 1 (**Fig 1D**). Intriguingly, *Hp* infection in mice fed high-fat diet exacerbated the frequency of the lymphoid aggregates, which were present in 14 of 17 mice, increasing the median limiting ridge inflammation score to 2. Taken together, these results demonstrate that *Hp* infection of mice fed a high-fat diet does not cause global gastric changes, but does increase inflammation at the limiting ridge.

### High-Fat Diet Impacts Metabolic and Defense-Related Pathways in the Stomach

We were surprised to observe no overt inflammation or substantial changes to the gastric tissue after ≥27 weeks on high-fat diet, given obesity’s known association with gastric disease ^16^. Therefore, we performed bulk RNA-sequencing (see **Supplementary Methods** for a detailed description of the analyses) using genetic material extracted from one-third portions of the mouse stomach (encompassing corpus and antrum) to explore biological pathways and processes that might change in response to diet-induced obesity and *Hp* infection. Principal component analysis (PCA) revealed that samples from mock-infected mice clustered separately from *Hp*-infected mice on PC2 (**Fig 2A**). However, within these two groups, mice did not cluster well based on diet. Among mock-infected mice, three mice in the high-fat group clustered with a standard diet mouse, while three mice in the standard diet group clustered with a high-fat diet mouse. Accordingly, when we compared differential gene expression between these two groups, there were no significantly differentially expressed genes (DEGs) meeting a false discovery rate (FDR) cutoff of 0.05 or a less stringent cutoff of 0.10. Therefore, we omitted the two outlier mice and compared the DEGs between the three remaining mock-infected, high-fat diet-fed mice vs. the three remaining mock-infected, standard diet-fed mice. Removing the two outliers yielded 776 DEGs (FDR<0.05), of which 25.5% were upregulated in the high-fat diet group (**Fig S2A, Table S1**). Among the most upregulated genes were leptin (*Lep*), a hormone that regulates long-term energy balance and fat stores in the body, and the lipase-related protein *Pnliprp2*. Several mitochondrial-related genes were also significantly upregulated, suggesting that high-fat diet drives alterations to energy metabolism. Conversely, several of the most downregulated genes, such as the defensins *Defa24* and *Defa30*, mannose-binding lectin *Mbl2*, and the regenerating genes *Reg1* and *Reg4*, are involved in antimicrobial defense, suggesting that high-fat diet may diminish gastric protection from infection. We used Ingenuity Pathway Analysis to evaluate the metabolic and signaling pathways impacted by high-fat diet (**Fig 2B, Table S2**). Only eight pathways met the significance cut-off (Benjamini-Hochberg *P* < 0.05), seven of which were downregulated in the high-fat diet-fed mice. Several pathways pertained to metabolism/digestion and nutrient processing, such as fructose metabolism, pancreatic secretion signaling, and sucrose degradation. Thus, high-fat diet has a modest impact on energy utilization and host defense in the stomach.

**Figure 2.**
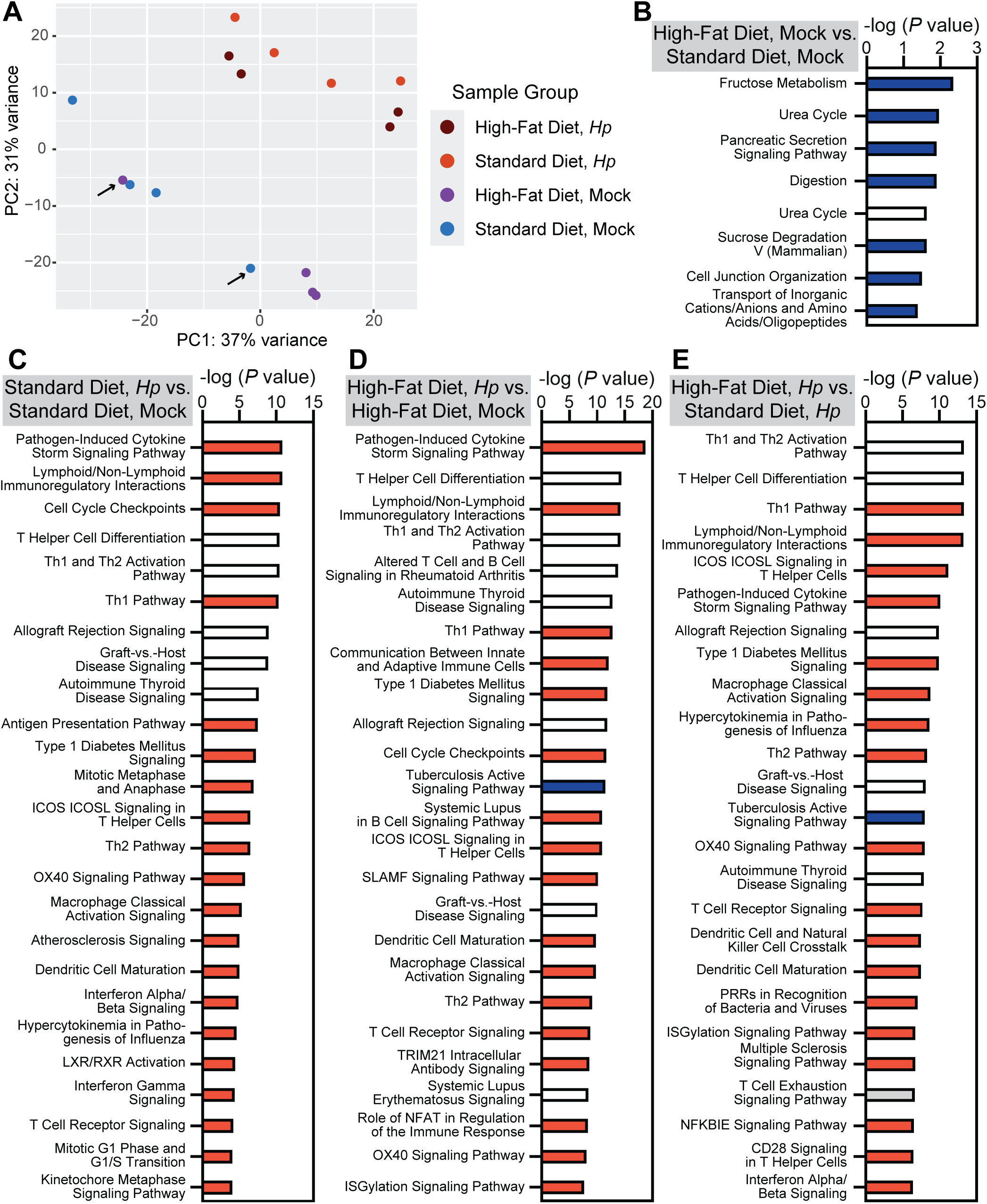
*Hp* infection elicits a robust inflammatory gene expression signature in the stomach, whereas high-fat diet has only a modest impact. Bulk RNA-sequencing was used to characterize gene expression in stomach tissues from C57BL/6 mice fed high-fat diet or standard diet and infected with *Hp* or mock-infected. Tissues were sequenced from n=4 samples per group originating from N=2 independent mouse experiments. **A)** In a principal component (PC) analysis, *Hp*-infected mice clustered separately from mock-infected mice. Arrows indicate two outlier mice from the mock-infected groups that were excluded from further analyses. **B-E)** Ingenuity Pathway Analysis results for the indicated comparisons are shown. Color-coding indicates the direction of the Z-score, which predicts up or downregulation of the pathway. Red bars show upregulated pathways (positive Z scores); blue bars show downregulated pathways (negative Z scores); the grey bar shows a pathway with a Z score of 0; white bars indicate pathways for which a Z score could not be determined. Z-score values are found in Table S2. All *P* values shown were adjusted for multiple comparisons. **B)** All statistically significant pathways are shown. Urea Cycle appears twice; the first (blue) pathway was curated by Reactome and the second (white) was curated by IPA. **C-E)** The top 25 most statistically significant pathways are shown.

### The Combination of Hp Infection and High-Fat Diet Increases Gastric Cldn7 and Reg3g Expression

In mice fed a standard diet, comparing *Hp* infection vs. mock infection yielded 3,133 DEGs, of which 54.9% were upregulated in the infected group (**Fig S2B, Table S1**). Among the most upregulated genes were several immunoglobulin genes, the T-cell co-receptor *Cd8b1*, and the complement-related genes *C9* and *Cfhr1*. Accordingly, pathway analysis revealed significant activation of immune-related pathways, such as pathogen-induced cytokine storm signaling, Th1 and Th2 pathways, and macrophage classical activation (**Fig 2C, Table S2**). For the comparison of *Hp* infection vs. mock infection in mice fed a high-fat diet, a similar phenomenon was observed. Out of 3,250 DEGs (59.1% of which were upregulated in the infected mice, **Fig S2C, Table S1**), some of the most significantly differentially expressed were numerous immunoglobulin genes, *Cd8b1*, and the interferon *Ifng*, suggesting enhanced B cell activation and antibody production as well as T cell activation. Overall, many of the DEGs and enriched pathways (**Fig 2D, Table S2**) were shared in both these comparisons, demonstrating that *Hp* infection activates similar inflammatory pathways and processes in the stomach whether or not diet-induced obesity is at play.

To investigate whether dietary state alters the gastric transcriptional response to *Hp* infection, we first assessed differential gene expression between *Hp*-infected mice fed high-fat diet vs. standard diet. There were 3,447 DEGs, of which 52.1% were upregulated in mice fed high-fat diet (**Fig S2D, Table S1**). However, the vast majority of the differentially expressed genes and pathways (**Fig 2E**) in this comparison were also differentially expressed in the comparisons of *Hp* infection vs. mock-infection with standard diet (**Fig 2C**) and with high-fat diet (**Fig 2D**). For example, all of the top 25 most significantly differentially expressed pathways (**Fig 2E, Table S2**) were also significantly differentially expressed in the other comparisons, though a few pathways, such as crosstalk between dendritic cells and natural killer cells and role of pattern recognition receptors in recognition of bacteria and viruses, fell outside the top 25 pathways. Thus, high-fat diet generally did not change <u>which</u> genes were induced by infection, but did have a modest impact on the <u>magnitude</u> of the gene expression.

Therefore, we investigated interaction effects using a generalized linear model with a diet x infection interaction term (see **Supplemental Methods**). We found that 44 genes were significantly differentially expressed (FDR<0.05), of which 28 showed a stronger response to the high-fat diet and 16 to the standard diet (**Fig S3A**). We used qRT-PCR to confirm the expression patterns of genes of interest in additional stomach samples (excluding any mice used for RNA-seq), this time using RNA extracted specifically from a 2-millimeter corpus biopsy punch. While the interaction was not reproducible for several genes (**Fig S3B**), possibly due to differences in sample type and/or heterogeneity among individual mice, the diet—infection interaction was reproducible for the tight junction gene *Cldn7* (**Fig 3A**) and the regenerating gene *Reg3g* (**Fig 3B**), two genes associated with gastric disease^46–48^. We performed immunostaining to assess CLDN7 localization in the stomach (**Fig 3C**); REG3G was not assessed due to lack of a reliable antibody. In mock-infected mice, regardless of diet, positive cells were primarily located at the base of the corpus glands. In *Hp*-infected mice fed a standard diet, a few brighter cells were observed higher in the glands. In *Hp*-infected mice fed a high-fat diet, staining was brighter at the base of the glands and extended higher up toward the gastric lumen, and more individual positive cells could be seen higher in the glands, supporting the increased gene expression in this group. Finally, we assessed *Hp* bacterial loads in the stomach via expression of the *Hp* 16S rRNA gene (**Fig 3D**). Interestingly, colonization was higher in the high-fat diet group (median relative gene expression level of 166) than in the standard diet group (median relative gene expression level of 22), though the difference did not reach statistical significance (*P* = 0.16).

**Figure 3.**
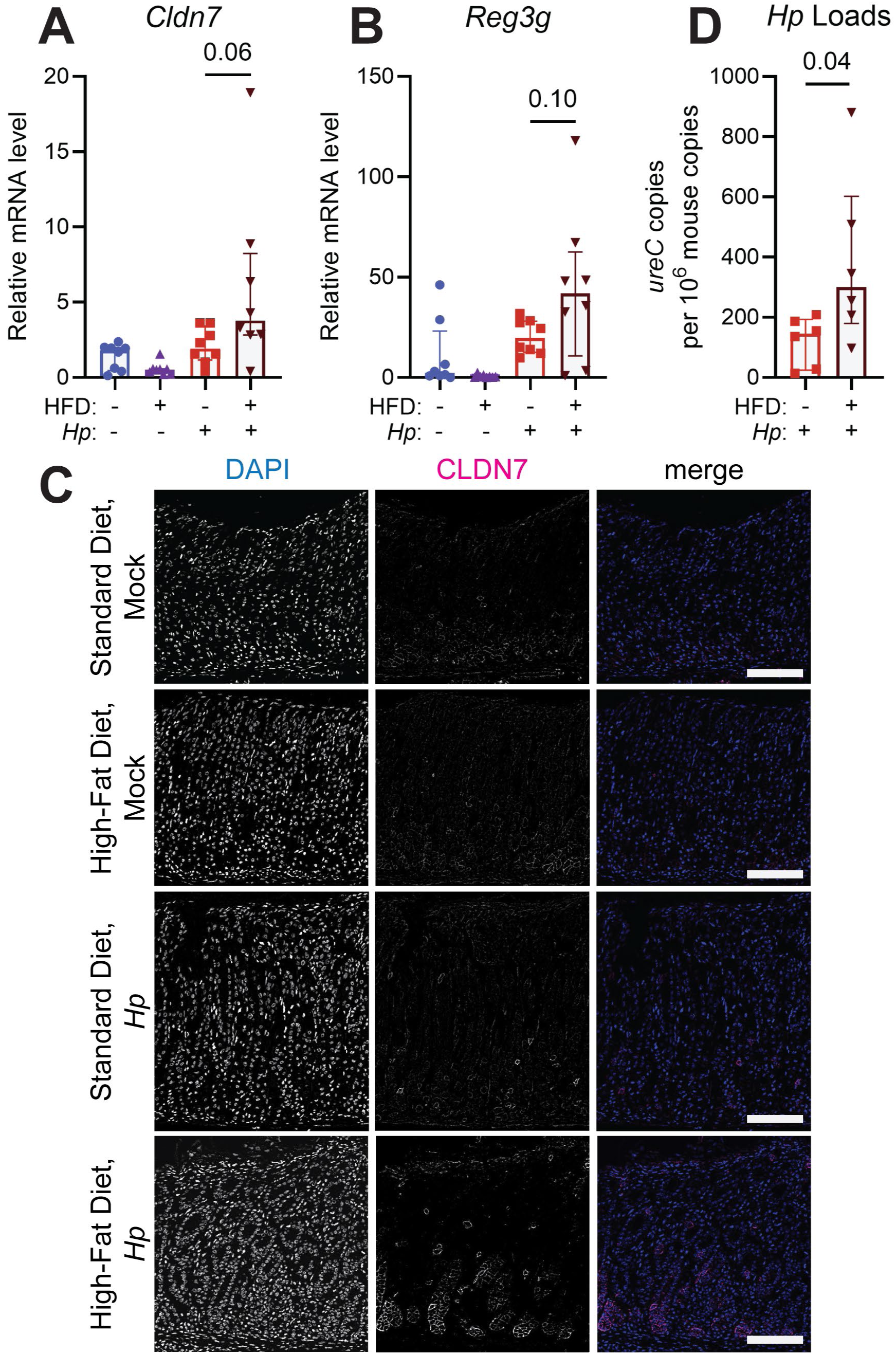
High-fat diet exacerbates gastric expression of *Cldn7* and *Reg3g* during *Hp* infection. Mice were fed standard diet or high-fat diet (HFD), and infected with *Hp* or mock-infected for 12 weeks. **A-B)** Expression of the indicated metaplasia-associated genes was assessed using qRT-PCR of RNA extracted from gastric corpus biopsy punches. Data are normalized to the expression level in mock-infected mice fed a standard diet (*i.e.*, healthy mice). **C)** Immunofluorescence microscopy was used to detect gastric corpus expression of CLDN7 (magenta in merge); DAPI (blue in merge) was used to stain nuclei. Representative images are shown. Scale bars, 100 µm. **D)** *Hp* stomach loads were assessed by qPCR to detect the relative DNA copy number of the *Hp ureC* gene relative to mouse housekeeping gene copies, x 10^6^. Statistical significance was assessed using the Mann-Whitney U test. Dots represent individual values for each mouse, bars indicate the median and error bars indicate the interquartile range. Data are from n=13-18 mice per group assessed in N=2 independent mouse experiments; n=6-8 mice per group were chosen randomly for qRT-PCR and qPCR.

### High-Fat Diet Exacerbates Metaplasia Phenotypes in Transgenic Mice

In wild-type mouse strains like C57BL/6 used above, 12 weeks of *Hp* infection does not elicit severe disease phenotypes ^44^. Here we found that incorporating high-fat diet still did not elicit frank metaplasia, but did reduce gastric expression of defense genes and induce the expression of the gastric disease-associated markers *Cldn7* and *Reg3g*. Therefore, we reasoned that high-fat diet might lower the threshold for pathological remodeling in the metaplastic setting. To test this hypothesis, we used the *Mist1-Kras* transgenic mouse model of gastric pseudopyloric and intestinal metaplasia. These mice have a tamoxifen-inducible Cre recombinase under the control of the *Mist1* promoter. Subcutaneous tamoxifen induces KRAS^G12D^ in *Mist1*-expressing cells, which in the stomach are progenitors and chief cells ^42, 43^. After six weeks, these “KRAS+ mice” exhibit preneoplastic phenotypes seen in humans including pseudopyloric metaplasia and mild dysplasia ^42^. We previously found that concomitant *Hp* infection (“*Hp*+KRAS+ mice”) significantly exacerbated disease in this model ^44^, with enhanced inflammation and the expansion of metaplastic pit cells that express the intestinal mucin *Muc4* ^45^. Here, after weaning, male and female *Mist1-Kras* mice were randomly assigned to receive high-fat diet or matched standard diet. After 15 weeks to induce obesity as indicated by body weight (**Fig S1B**), mice were infected with *Hp* or mock-infected, and KRAS^G12D^ was induced with subcutaneous tamoxifen. Mice were euthanized six weeks later and gastric disease was assessed by pathology scoring, qRT-PCR, and *in situ* hybridization.

We first performed a blinded analysis (**Fig 4A-B**) and scoring (**Fig 4C-G**) of tissue pathology using a modified version of the Rogers histological activity index (see **Supplementary Methods** and **Fig S4**). Disease pathology was exacerbated by both *Hp* infection and high-fat diet (**Fig 4A-B**). We found that compared to standard diet, high-fat diet increased the severity of several preneoplasia phenotypes. **Fig 4C-F** shows the proportion of mice in each group with the indicated score, whereas **Fig S4** shows the individual scores for each mouse. High-fat diet caused increased hyperplasia (**Fig 4C**), pseudopyloric metaplasia (**Fig 4D**), and dysplasia (**Fig 4E**) in KRAS+ mice, with a greater proportion of high-fat diet-fed mice scoring 4 for hyperplasia (complete loss of functional mucosa), 4 for pseudopyloric metaplasia (replacement of >90% of the oxyntic mucosa by antralized glands), and 1 for dysplasia (presence of occasional multifocal dysplastic glands marked by elongation, altered shapes, and back-to-back forms). *Hp*+KRAS+ mice fed a standard diet scored similarly to mock-infected mice fed a high-fat diet, indicating that both high-fat diet and *Hp* infection augmented these disease phenotypes to a similar extent. *Hp*+KRAS+ mice fed a high-fat diet were the most likely to score 4 for hyperplasia and pseudopyloric metaplasia and 1 for dysplasia, with an increased number of deep glands with the dysplastic characteristics of disorganized and piled-up cells, increased branching, loss of polarity, and nuclear and cellular pleomorphism (**Fig 4B**). The pattern for oxyntic atrophy was slightly different (**Fig 4F**); while high-fat diet did increase the proportion of mice scoring 4 (no chief cells, and few/no parietal cells), *Hp* infection had a greater impact, and all *Hp*+KRAS+ mice fed high-fat diet scored a 4. There were no differences in inflammation or epithelial defects (**Fig S4**). The overall histology score sums these individual parameters.

**Figure 4.**
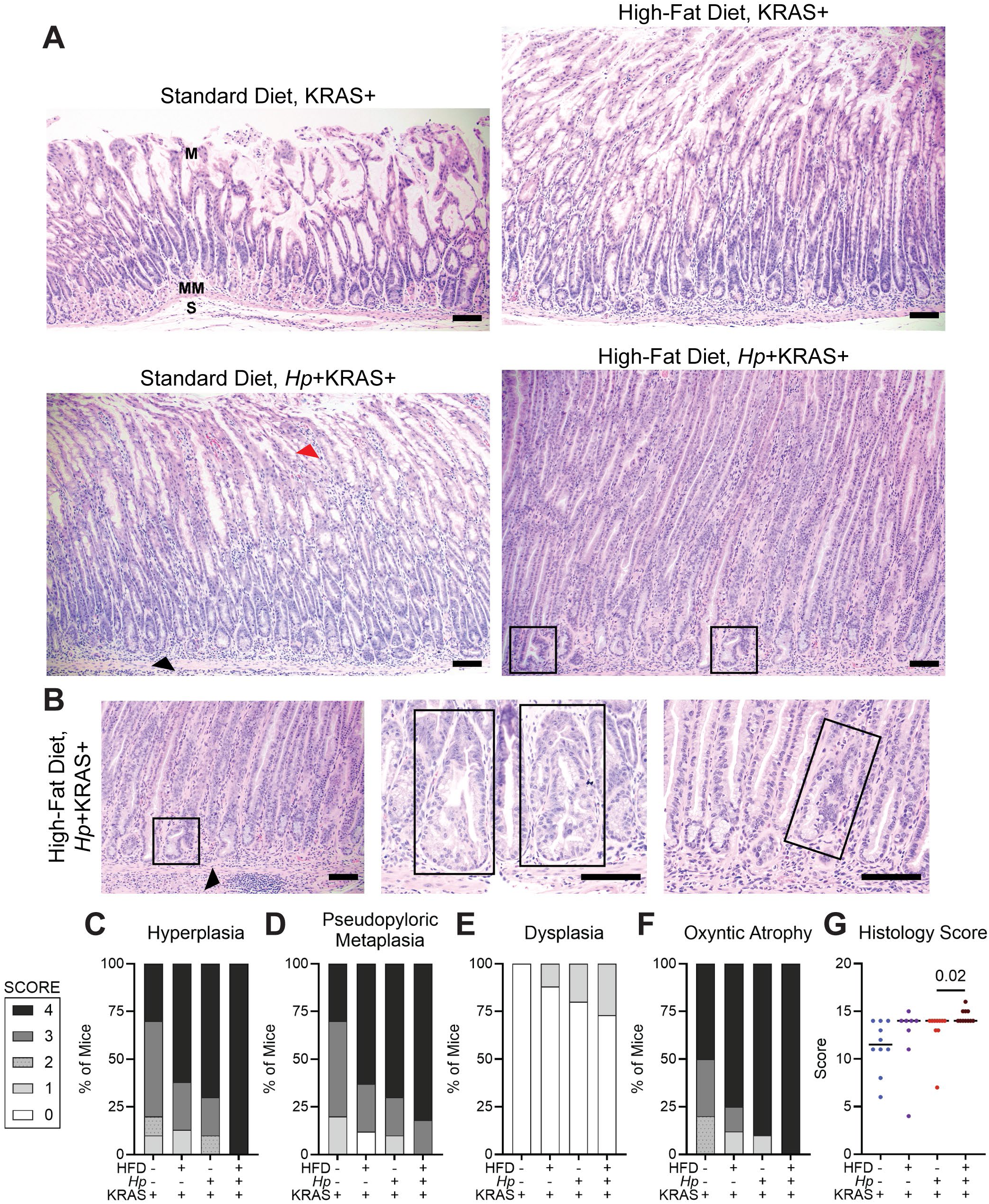
Histopathology scores are highest in *Hp*+KRAS+ mice fed a high-fat diet. Mice with obesity induced by high-fat diet (HFD), or control mice fed a standard diet, were infected with *Hp* or mock-infected, then tamoxifen was used to induce KRAS^G12D^. After six weeks, mice were euthanized and a blinded analysis of tissue pathology was performed. **A-B)** Representative images are shown. Scale bars, 100 µm. **A)** The low-power field images show the representative tissue pathology of each of the four treatment groups. M, mucosa; MM, muscularis mucosa; S, submucosa. Red arrowhead shows an example of inflammatory cells higher up in the mucosa and black arrowhead shows an example of inflammatory cells deep in the mucosa. Boxes indicate examples of dysplastic glands. **B)** Additional images from *Hp*+KRAS+ mice fed high-fat diet are shown, with boxes indicating examples of dysplastic glands. The left, low-power field image shows one of the same dysplastic glands seen in **A** along with a lymphoid aggregate in the submucosa (black arrow). The middle and right, high-power field images show additional dysplastic glands. **C-F)** The indicated parameters were scored from 0 to 4 (see color key left of panel **C**). Data are represented as the proportion of mice exhibiting each score; individual scores for each mouse are given in Figure S4. **G)** The total histology score is shown for each mouse. Statistical significance was assessed using the Mann-Whitney U test. Dots represent individual values for each mouse and bars indicate the median. Data are from n=8-11 mice per group assessed in N=2 independent mouse experiments.

The median was 11.5 for *Hp*-KRAS+ mice fed standard diet, our baseline group, and 14 in each of the other three groups (**Fig 4G**). Although the median was the same in *Hp*+KRAS+ mice fed standard vs. high-fat diet, the high-fat diet group had uniformly high scores, whereas the standard diet group had three lower-scoring mice. Consequently, the overall histology score was statistically significantly increased in *Hp*+KRAS+ mice fed high-fat diet (*P* = 0.02). Together, the pathology scoring of both the C57BL/6 mice and transgenic mice suggests that while high-fat diet does not directly cause preneoplasia, it can exacerbate certain preneoplastic phenotypes.

### High-Fat Diet Alters Metaplasia Marker Expression

We used qRT-PCR to assess gene expression changes in corpus biopsy samples from the transgenic mice, compared to healthy control mice. In accordance with its association with gastric cancer ^46, 47^, *Cldn7* expression was upregulated in all metaplasia groups compared to healthy control mice. While high-fat diet caused a significant increase of *Cldn7* expression in KRAS+ mice, expression was greater in *Hp*+KRAS+ mice and did not further increase with diet treatment (**Fig S5**). *Reg3g* expression was minimally induced by metaplasia alone; expression increased with either high-fat diet or *Hp* infection, but again, there was no statistically significant difference between *Hp*+KRAS+ mice fed standard diet vs. high-fat diet (**Fig S5**). We evaluated CD44v10 (the mouse ortholog of human CD44v9) as a marker of pseudopyloric metaplasia^49^. High-fat diet increased CD44v10 gene and protein expression in KRAS+ mice (**Fig S6**). We next assessed mucin expression patterns, because metaplasia is well known to alter the mucin profile in the stomach. The intestinal metaplasia marker *Muc2* was not highly expressed in any group (**Fig 5A**), in line with previous findings for mice fed a standard diet ^44^. *Muc6*, which is expressed in mucous neck cells and pseudopyloric metaplasia, increased in KRAS+ mice fed a high-fat diet compared to standard diet (*P* = 0.08, **Fig 5B**), in accordance with the increased expression of *CD44v10*. In *Hp*+KRAS+ mice, the effect was the opposite, with *Muc6* significantly decreasing in the high-fat diet group compared to standard diet (*P* = 0.03). *Muc13*, a mucin associated with *Helicobacter*-driven metaplasia ^50^, had heterogeneous expression in all groups (**Fig 5C**). The median increased in each of the high-fat diet groups compared to their standard diet counterparts, but the differences were not statistically significant. Finally, *Muc4*, which we previously identified as a marker for inflammation-associated metaplastic pit cells ^45^, increased in KRAS+ mice fed high-fat diet (*P* = 0.07, **Fig 5D**), and significantly increased in *Hp*+KRAS+ mice fed high-fat diet (*P* = 0.04). The latter group had the highest mean and median *Muc4* expression.

**Figure 5.**
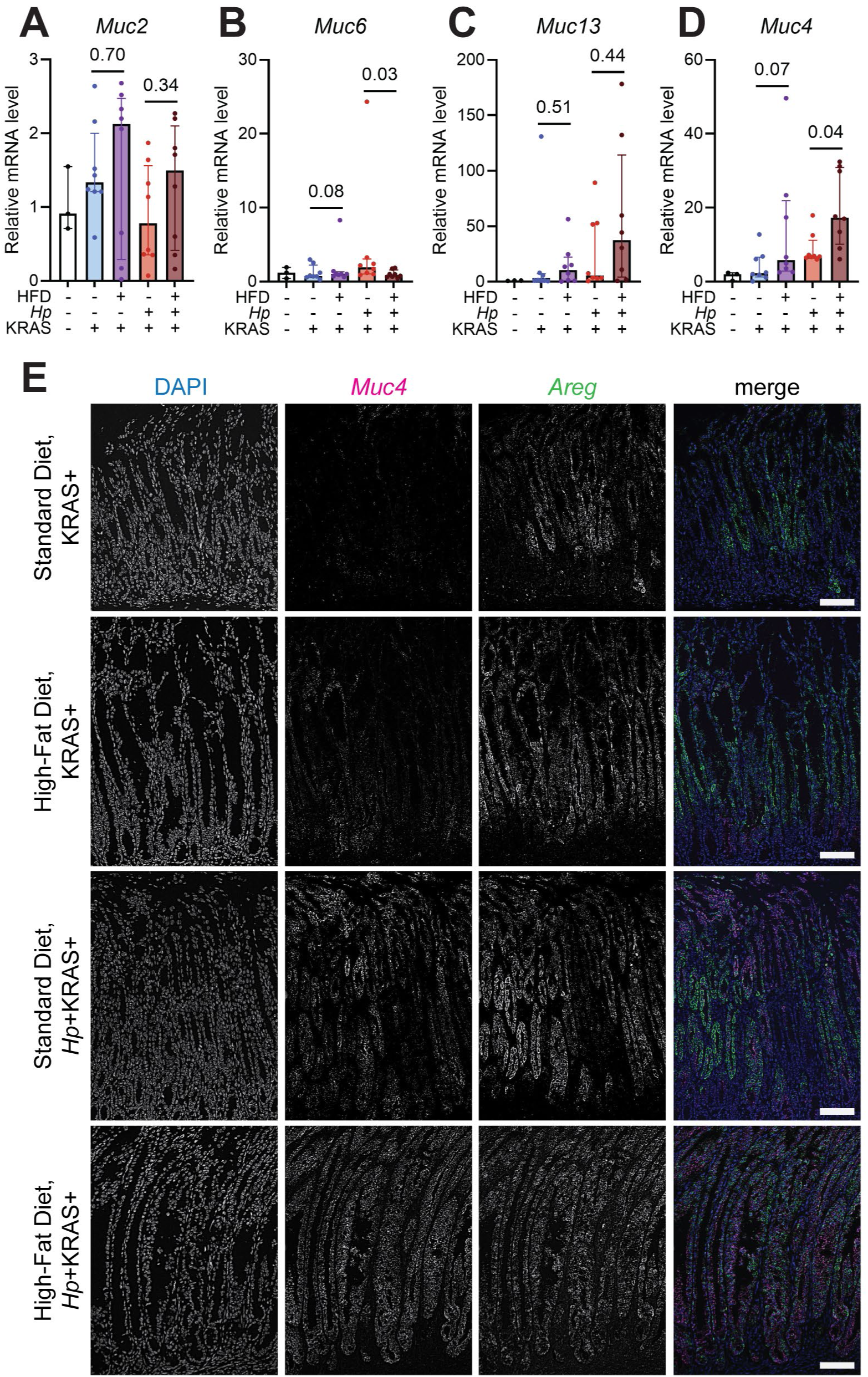
High-fat diet promotes metaplastic pit cell expansion during metaplasia. **A-D)** Expression of the indicated metaplasia-associated mucins was assessed using qRT-PCR of RNA extracted from gastric corpus biopsy punches, relative to corpus biopsy RNA from n=3 healthy control mice (white bars, black dots). Statistical significance was assessed using the Mann-Whitney U test. Dots represent individual values for each mouse, bars indicate the median and error bars indicate the interquartile range. Data are from n=8-11 mice per treatment group assessed in N=2 independent mouse experiments; n=8 mice per group were chosen randomly for qRT-PCR. **E)** *In situ* hybridization was used to detect gastric corpus expression of the metaplastic pit cell markers *Muc4* (magenta in merge) and *Areg* (green in merge). Representative images are shown. Scale bars, 100 µm.

To further characterize metaplastic pit cells, we performed *in situ* hybridization to detect the metaplastic pit cell markers *Muc4* and amphiregulin (*Areg*) in gastric corpus tissue sections (**Fig 5E**). In accordance with the qRT-PCR results, *Muc4* expression was sparse in KRAS+ mice fed a standard diet, and was mostly seen at the base of the glands. *Areg*, which is also expressed by other gastric cell types such as parietal cells ^51^, was more evident in these tissues, but still patchy. The addition of high-fat diet increased *Muc4* expression in KRAS+ mice, including in cells located higher up in the glands (toward the gastric lumen), and reduced the patchiness of *Areg* expression. In *Hp*+KRAS+ mice fed a standard diet, *Muc4* was widespread throughout the corpus, with most glands co-expressing *Muc4* and *Areg*. The addition of high-fat diet in *Hp*+KRAS+ mice further expanded the metaplastic pit cell population, with robust *Muc4* and *Areg* staining throughout the corpus. Together, these results demonstrate that high-fat diet increases the expression of genes associated with pseudopyloric metaplasia and metaplastic pit cells in the stomach.

## DISCUSSION

*Hp* infection is the biggest risk factor for gastric cancer development, with about 80% of gastric cancers attributed to this bacterium ^1^. However, only 1-2% of infected individuals develop gastric cancer ^6, 52^. Therefore, additional risk factors beyond *Hp* infection tip the balance toward cancer development in some individuals. Known risk factors include tobacco and alcohol, host genetic background, and being male ^6, 53^. One risk factor associated with many types of cancer is obesity. Obesity is strongly associated with cardia cancer ^18, 19^. Intriguingly, we observed that the combination of high-fat diet and *Hp* infection significantly increased inflammation at the limiting ridge, a comparable anatomical site in mice. For non-cardia cancer, the more predominant type that occurs in the middle and lower portions of the stomach, the association with obesity is less clear. Some studies report that increased body mass is associated with greater cancer incidence and death ^20–24^, whereas other studies find no association ^54–56^. A key confounder is that metaplasia, a precancerous state that can appear many years before stomach cancer develops ^57^, frequently causes weight loss ^58^, so studies that assess body weight at time of cancer diagnosis or even years prior may fail to capture the true weight trajectories of individuals during the prolonged inflammatory period that precedes metaplasia. Therefore, we sought to test the hypothesis that diet-induced obesity could increase gastric cancer risk by augmenting *Hp*-driven preneoplastic disease phenotypes. We found that in a transgenic model in which metaplasia is driven by induction of constitutively active KRAS^G12D^ and exacerbated by *Hp* infection ^42, 44, 45^, the addition of obesity induced by a diet very high in fat (60% kilocalories from fat) worsened preneoplastic phenotypes, causing increased oxyntic atrophy, hyperplasia, pseudopyloric metaplasia, and dysplasia. Diet-induced obesity augmented disease pathology in both KRAS+ mice and *Hp*+KRAS+ mice, with the most severe gastric pathology seen in *Hp*+KRAS+ mice fed high-fat diet. However, in C57BL/6 mice, diet-induced obesity had a very modest effect on the tissue; we observed minimal diet-dependent gastric pathology and relatively few gene expression changes. Thus, in our pre-clinical models, obesity was not sufficient to elicit metaplasia, but did exacerbate genetically-driven preneoplastic phenotypes.

In a previous study, C57BL/6 mice were infected with *Helicobacter felis*, a related species that causes enhanced stomach inflammation in mice compared to *Hp* ^59^, and then high-fat diet (45% kilocalories from fat) was initiated three weeks later. Mice remained on the diet for ten to 15 months, at which time points metaplasia and dysplasia were significantly increased in infected mice fed high-fat diet vs. standard diet ^60^. Inflammation was also increased, with a significant increase in CD4 T cells and gastric IL-17A production suggesting a T_H_17 response. In our study, C57BL/6 mice were infected for only three months; upregulation of gastric *Cldn7* and *Reg3g* was among the only unique change we observed in *Hp*-infected mice fed a high-fat diet. Extending our studies to ten to 15 months could reveal more substantial immunopathology in *Hp*-infected mice fed high-fat diet. Alternatively, the phenotypes observed in the prior study may be unique to the *H. felis*—high-fat diet interaction. Nonetheless, the genes we identified are of interest because of their specific functions and known associations with gastric disease. Claudin-7 is a tight junction protein that is highly expressed in the intestinal epithelium; claudin-7-deficient mice die rapidly after birth from severe inflammatory bowel disease ^61^. In human studies, claudin-7 was upregulated in metaplasia and gastric cancer samples compared to healthy control gastric samples ^46, 47^. Although increased *Cldn7* expression in C57BL/6 mice is consistent with changes reported in metaplastic stomach tissue, the absence of histologic metaplasia in C57BL/6 mice suggests that *Cldn7* expression is not an indicator of metaplasia. Instead, it may reflect an early intestinalization program that precedes overt metaplastic transformation, and/or pathological barrier remodeling induced by *Hp* infection in the context of high-fat diet. Reg3g is an antimicrobial peptide (with activity primarily against Gram-positive bacteria; *Hp* is Gram-negative) that is induced in the intestinal tract by bacterial colonization ^48^. One study found that REG3γ was overexpressed in biopsy samples from human subjects with *Hp* gastritis compared to disease-free control tissues; expression was dependent on the presence of the *Hp cag* type IV secretion system effector CagA ^62^. The *Hp* strain used in our studies, PMSS1, encodes an active *cag* secretion system and the CagA toxin ^63^. In our RNA-seq experiment, we observed decreased expression of antimicrobial defense genes in mock-infected C57BL/6 mice fed high-fat diet. Thus, diet-induced obesity may allow *Hp* and/or other members of the gastric microbiota to colonize the stomach to greater levels. Indeed, we observed increased *Hp* loads in mice fed high-fat diet compared to standard diet. Increased bacterial colonization could be the trigger for increased *Reg3g* expression in our model. Together, the altered expression of epithelial defense and barrier genes we observed suggests that diet-induced obesity may place the gastric epithelium in a remodeled or stressed state that alters its response to infection. Future studies will dissect the mechanisms for *Cldn7* and *Reg3g* induction in *Hp*-infected mice fed high-fat diet.

Notably, in the aforementioned study ten to 15 months of high-fat diet did not cause gastric inflammation, metaplasia, or dysplasia in mock-infected mice ^60^, which is similar to our observations in mock-infected mice fed high-fat diet for about six months (15 weeks to induce obesity followed by a 12 week study period). Together, our work and the previous study strongly suggest that high-fat diet alone does not cause gastric immunopathology. In contrast, another group previously reported that 12 weeks of the same high-fat diet we used, D12492i from Research Diets, induced stomach metaplasia in C57BL/6 mice ^64, 65^. Possibly, mice that reportedly developed metaplasia from a high-fat diet may have harbored metaplasia-inducing microbes that expanded in response to this dietary intervention and drove the disease.

Several studies have reported a high prevalence of gastritis (stomach inflammation) in obese subjects ^31, 32^. While our immunopathology scoring did not reveal significantly increased corpus inflammation among different mouse groups, we note that the system used to score inflammation in these mice is based primarily on lesion distribution rather than the absolute number of immune cells ^66^, and therefore, the numbers of immune cells may differ between mice with the same score. Interestingly, a short-term period of high-fat diet feeding (60% kilocalories from fat for three weeks) in male C57BL/6 weanlings led to expansion of gastric mast cells, as well as tuft cells and pit cells ^67^. It is possible that acute vs. chronic high-fat diet feeding could differentially impact gastric inflammation. The observation that pit cells expanded with short-term high-fat diet feeding also could suggest that pit cells are uniquely sensitive to high-fat diet. Accordingly, we found that *Muc4* expression was increased in *Hp*+KRAS+ mice fed a high-fat diet. We previously found that *Muc4*-expressing metaplastic pit cells were a hallmark of preneoplasia in *Hp*+KRAS+ mice and were associated with *Hp*-driven inflammation ^45^. Future studies will dissect whether the expanded metaplastic pit cells in high-fat diet-fed mice is due to modulated gastric inflammation vs. direct effects of the dietary intervention on pit cells or their progenitors.

A key limitation of our study is the timing of the dietary intervention vs. the infection. Humans are typically infected with *Hp* during childhood, whereas obesity typically develops later in life. Future studies in which mice are first infected with *Hp* and then fed a high-fat diet to induce obesity would more accurately reflect the natural course of disease. Notably, *Hp* infection of neonatal mice was shown to elicit a tolerogenic response that protected against severe gastric disease, demonstrating that host age at time of infection significantly impacts the gastric immune response ^63^; whether subsequent high-fat diet feeding could break this tolerance would be of interest to test. In contrast, our study tests the specific hypothesis that host-pathogen interactions may differ in the context of diet-induced obesity, which is relevant to the overarching question of whether and how obesity may promote gastric cancer development. As well, we induced obesity in the transgenic *Mist1-Kras* mice prior to inducing metaplasia via tamoxifen administration; this sequence of events likely reflects the natural course of disease in humans. We note that *Mist1-Kras* mice model metaplasia, not cancer. Specifically, these mice exhibit pseudopyloric metaplasia and metaplastic pit cells (both phenotypes increased by *Hp* infection) as well as rare, MUC2-expressing goblet cells (not increased by *Hp* infection) indicative of intestinal metaplasia ^44, 45, 68^. A previous study found that murine forestomach carcinoma cells grew faster when injected into the flanks of mice with diet-induced obesity than in lean mice ^69^; however, the forestomach is a squamous epithelium more similar to the esophagus than the glandular stomach, so whether obesity impacts gastric cancer development in mice is still an open question.

Another limitation of our study is the composition of the dietary intervention: 60% kilocalories from fat, primarily lard. First, we note that the phenotypes we observed may be due to direct gastric exposure to fatty acids, rather than systemic inflammatory and/or metabolic changes associated with obesity. Second, a typical Western diet provides only 36-40% of its calories from fat ^70^. Nonetheless, researchers commonly use the 60% fat diet because it rapidly induces obesity, allowing for shorter studies, and phenotypes in these mice are typically only modestly more severe than in mice with obesity induced by diet with 45% kilocalories from fat ^70^. However, it is possible that obesity induced by a lower-fat diet or a diet higher in sucrose would have a different impact on host-pathogen interactions in the stomach. Likewise, future studies investigating *Hp* infection in mice with genetically-driven obesity (*e.g.*, ob/ob or db/db mice) would help clarify whether our phenotypes were driven by obesity or fatty acid exposure.

In conclusion, while *Hp* infection is the primary risk factor for gastric cancer development, obesity is another factor that has been shown to exacerbate gastric cancer risk ^20–24^. While most human studies about obesity and gastric cancer have not evaluated *Hp* infection status, one study did find that obesity and *Hp* were independent risk factors for gastric cancer development ^20^. Likewise, obesity is associated with stomach metaplasia development, independent of *Hp* infection status ^17^. We found that approximately six months of high-fat diet alone in C57BL/6 mice did not induce gastric inflammation or precancerous changes, but high-fat diet did augment gastric *Hp* infection in these mice, leading to increased *Hp* stomach loads after 12 weeks of infection and increased gastric *Cldn7* and *Reg3g* expression. In contrast, in the *Mist1-Kras* transgenic model of stomach metaplasia, high-fat diet exacerbated preneoplastic phenotypes in both KRAS+ mice and *Hp*+KRAS+ mice, with increased hyperplasia, pseudopyloric metaplasia and metaplastic pit cell marker expression, and dysplasia. Collectively, these findings suggest that diet-induced obesity may alter epithelial defenses and barrier-associated programs in the stomach, creating a permissive environment for microbial-driven tissue remodeling and metaplastic progression under susceptible conditions.

## MATERIALS AND METHODS

Additional information is found in the Supplementary Materials and Methods section of the supporting information.

All animal studies were approved by the Purdue University Institutional Animal Care and Use Committee (protocol 523002388). C57BL/6J and *Mist1-Kras* ^42^ (*Mist1-CreERT2^Tg^*^/+^; *LSL-K-Ras*(*G12D*)*^Tg^*^/+^) mice were maintained under specific pathogen-free conditions and randomly assigned to either a control standard diet comprising 10% kilocalories from fat (Research Diets, Inc; D12450J) or a high-fat diet comprising 60% kilocalories from fat (Research Diets, Inc; D12492). Fifteen weeks on high-fat diet was sufficient to induce obesity in males and females based on body weight. After 15 weeks, mice were infected with the *Helicobacter pylori* clinical isolate PMSS1 (10^8^ CFU) or mock-infected with culture medium, via oral gavage. *Mist1-Kras* mice received tamoxifen (5 mg/day for 3 consecutive days) beginning one day after infection to induce KRAS^G12D^ ^44^. C57BL/6J mice were euthanized 12 weeks post-infection and *Mist1-Kras* mice six weeks post- infection. Two independent experiments were performed in each mouse line, with final group sizes of 13-18 C57BL/6J mice and 8-11 *Mist1-Kras* mice per treatment.

After euthanasia, stomachs were harvested and divided into portions for histologic, immunofluorescence, *in situ* hybridization, and gene expression analyses. Formalin-fixed, paraffin-embedded sections were stained for CLDN7 (Life Technologies, 34-9100) and imaged using a Keyence BZ-X800 microscope. Expression of *Muc4* and *Areg* transcripts was assessed in fixed sections by RNAscope (ACD Biotechne) according to the manufacturer’s instructions. Hematoxylin and eosin-stained sections were evaluated in a blinded manner by a board-certified veterinary pathologist (VVP). C57BL/6J tissues were scored for inflammation, while *Mist1-Kras* tissues were scored for preneoplasia phenotypes using a previously established scoring system ^66^.

Total DNA or RNA was isolated from glandular stomach tissue using TRIzol (ThermoFisher Scientific; 15596026). For qRT-PCR to detect gene expression changes, eight mice per group were randomly selected, cDNA was prepared and gene expression was quantified using SYBR Green (Applied Biosystems, A46112) on a QuantStudio 6 Real-Time PCR system (Applied Biosystems, 4485691), and normalized to the average of five housekeeping genes (*Hprt, Ppia, H2afz, Actb,* and *Tbp*) using the ΔΔCt method. For qPCR to detect *Hp* abundance in the stomach, *Hp ureC* copies and mouse housekeeping gene copies were detected using SYBR Green and analyzed using the ΔCt method. Primers are listed in the Supplementary Materials and Methods. RNA-seq was performed on stomach tissue from four C57BL/6J mice per treatment group by Novogene (Sacramento, USA). Libraries were sequenced on an Illumina NovaSeq X Plus platform. Reads were processed using fastp v0.23.2 ^71^ and STAR v2.7.11.b ^72^ was used to map data to the *Mus musculus* GRCm39 reference genome. FeatureCounts ^73^ was used to count reads mapping to genes and edgeR v.4.0.16 ^74, 75^ was used within R v4.3.3 to perform statistical analyses. Differentially expressed genes were defined as FDR < 0.05, and pathway analyses were performed using Ingenuity Pathway Analysis (Qiagen). Other statistical analyses were conducted using Mann-Whitney U tests in GraphPad Prism v10.5.0. RNA-seq data have been deposited at GEO and will be released upon publication; all other data, metadata, and methods used to reach the conclusions in the paper and any additional data required to replicate the study findings will be provided upon request.

## ACKNOWLEDGEMENTS

The authors wish to thank Anne Dewar and Margaret Bright for technical assistance. This work was supported by the National Cancer Institute (R00CA263036 to VPO) and the V Foundation for Cancer Research (V2025-030 to VPO). As well, the authors gratefully acknowledge the support of the Purdue Institute for Cancer Research (NIH grant P30CA023168), the Walther Cancer Foundation, and the Jim and Diann Robbers Cancer Research Grant for New Investigators Award to VPO. The funders had no role in study design, data collection and interpretation, or the decision to submit the work for publication.

## Notes

### Competing Interest Statement

The authors have declared no competing interest.

### Summary of Updates

Section on bacterial burdens updated to a qPCR assay detecting Hp DNA copies relative to host DNA copies; pathology images annotated and additional images of dysplasia shown; gene and protein expression of the metaplasia marker CD44v10 analyzed.

