## Supplemental Methods and Figures for "Diet-Induced Obesity Exacerbates *Helicobacter pylori*-Associated Precancerous Phenotypes"

**Supplemental Material Table of Contents**

Supplemental Methods, pages 2-6

Supplemental Figures, pages 7-13

Supplemental Excel Tables containing RNA-seq data: uploaded separately

**Supplemental Methods**

Mouse experiments

All mouse experiments were approved by the Purdue University Institutional Animal Care and Use Committee (IACUC), protocol number 523002388. Male and female mice (specific pathogen-free) were housed in cages docked in HEPA-filtered ventilation racks with airflow control. Mice were kept on a 12 hour light/dark cycle and had access to chow and water ad libitum. Mice were housed in groups of no more than 5 per cage for standard diet and no more than 4 per cage for high-fat diet. C57BL/6J mice (bred in a clean facility in the Hansen Life Sciences Research Building at Purdue University, using breeder mice from the Jackson Laboratory) were randomly assigned to a control standard diet comprising 10% kilocalories from fat (Research Diets, Inc; D12450J) or a high-fat diet comprising 60% kilocalories from fat (Research Diets, Inc; D12492) at the time of weaning. After 15 weeks to induce obesity, cages were randomized to receive infection with $10^8$ colony-forming units of the *H. pylori* clinical isolate PMSS1 or mock (broth) via oral gavage. *Mist1-Kras* mice (*Mist1-CreERT2*$^{Tg/+}$; *LSL-K-Ras*(*G12D*)$^{Tg/+}$; C57BL/6 background) were previously described [27]. Mice were imported to Purdue University from Fred Hutchinson Cancer Center and re-derived as specific pathogen-free, then housed in the same clean facility in the Hansen Life Sciences Research Building. *Mist1-Kras* mice were randomly assigned to receive the same diets described above, and after 15 weeks to induce obesity, mice were transferred to an animal biosafety level 2 facility at Purdue University. After approximately one week to acclimate, mice were infected or mock-infected by oral gavage as above. Starting the next day, mice received three daily subcutaneous doses of 5 mg tamoxifen in corn oil as previously described [22]. C57BL/6 mice were euthanized 12 weeks after infection and *Mist1-Kras* mice were euthanized 6 weeks after infection, and stomachs were analyzed. Two independent replicates of each experiment were performed, with a total of n=13-18 C57BL/6 mice per group and n=8-11 *Mist1-Kras* mice per group.

*Helicobacter pylori* growth, infections, and stomach processing

The model *Hp* strain PMSS1 was stored at -80°C and recovered on *Hp* horse blood plates (4.4% Columbia blood agar base (Fisher, 279220), 10 µg/ml Vancomycin (GoldBio, V-200-5), 5 µg/ml Cefsulodin (GoldBio, C-114-5), 2.5 U/ml Polymyxin B sulfate (Fisher, P19231G), 5 µg/ml Trimethoprim (GoldBio, T-350-5), 8 µg/ml Amphotericin B (GoldBio, A-560-5), 0.2% Beta-cyclodextrin (Fisher, AAA1452922), 5% defibrinated horse blood (HemoStat, DSB100)) grown under microaerophilic conditions (10% $CO_2$, 10% $O_2$). *Hp* cells were then expanded in liquid culture in "BB10" media, comprising Brucella Broth (Fisher, B11088) with 10% heat-inactivated fetal bovine serum (R&D Systems, S11550H), grown shaking (200 rpm) in microaerophilic conditions. Mice were inoculated with $10^8$ colony-forming units of *Hp* in 100 µl of BB10, or 100 µl of BB10 as a control (mock), via oral gavage. After 6-12 weeks, mice were humanely euthanized via $CO_2$ inhalation and stomachs were removed aseptically for processing. Mice were not fasted prior to euthanasia. The stomach was opened along the lesser curvature and spread flat, and forceps were used to gently scrape off food. A scalpel blade was used to cut stomachs into one-third portions, each spanning corpus and antrum. One portion was frozen on dry ice, stored at -80°C, and used for RNA extraction for RNA-sequencing experiments. One portion was spread flat onto Whatman paper and fixed for 48 hours in 10% neutral-buffered formalin, then processed for thin sections as described below. From the final portion, a 2 mm corpus biopsy punch was collected and stored in TRIzol and used for RNA extraction for qRT-PCR. The rest of this portion was used for method optimization. Tissue sectioning and immunofluorescence imaging

The Histology Research Laboratory in the Purdue Center for Comparative Translational Research embedded formalin-fixed sections in paraffin blocks and cut thin (5 µm) sections. To assess CLDN7, CD44v10, and clusterin expression, paraffin-embedded tissues were deparaffinized using three changes of Histo-Clear (Fisher, 50-899-90147) and rehydrated using decreasing concentrations of ethanol. Antigen retrieval was performed in a pressure cooker for 15 minutes with 1:100 diluted Antigen Retrieval Buffer (Abcam, AB93684) followed by washing in water. Tissues were pretreated with a 1:10 dilution of hydrogen peroxide (Fisher, H325-500) in methanol (Fisher Chemical, A4521) for 10 minutes prior to autofluorescence quenching. After peroxide/methanol pretreatment, slides were washed in water and PBS and then submerged in PBS for autofluorescence quenching using the TiYO Autofluorescence Quenching system (NepaGene) for 60 minutes.

Tissues were incubated in Protein Block, Serum-Free (Agilent, X090930) for 90 minutes at room temperature. Rabbit anti-CLDN7 (Life Technologies, 34-9100) was diluted 1:100 in Antibody Diluent, Background-Reducing (Agilent, S302281-2) and slides were incubated overnight at 4°C. Rat anti-mouse CD44v10 (Cosmo BIO, CAC-LKG-M002) was diluted 1:100 in PBS followed by 1:250 into antibody diluent, while goat anti-CLU (Fisher, AF2747SP) was diluted 1:250 into antibody diluent. Tissues were washed in three changes of PBS  Secondary antibodies were diluted 1:500 in Protein Block, Serum Free and applied for 60 minutes at room temperature, protected from light. Secondaries used were: for CLDN7, Alexa Fluor 647 donkey anti-rabbit IgG; for CD44v10, Alexa Fluor 488 donkey anti-rat IgG; and for clusterin, Alexa Fluor 647 donkey anti-goat IgG (Invitrogen, A31573, A21208, A21447, respectively). Slides were washed in three changes PBS protected from light, followed by a dip in water. To visualize nuclei, 1:1000 DAPI in water (Fisher, EN63348) was applied to tissues for 10 minutes. Tissues were mounted with ProLong Diamond Antifade Mountant (Fisher, P36961). Slides were imaged on a Keyence BZ-X800 fluorescence microscope.

*In situ* hybridization

*Muc4* and *Areg* were visualized in thin tissue sections using the RNAscope Multiplex Fluorescent Detection Kit v2 (ACD Biotechne, 323110) according to the manufacturer's instructions. Briefly, slides were deparaffinized by incubating at 60°C for 60 minutes followed by two xylene and two ethanol incubations. Slides were dried at 60°C for 5 minutes and covered in hydrogen peroxide for 10 minutes. After a water wash, slides were placed in boiling antigen retrieval solution for 15 minutes. Slides were washed in water and ethanol, then dried at 60°C for 5 minutes. Protease Plus was applied to the tissues and incubated at 40°C for 15 minutes, followed by a water wash. To hybridize probes, *Muc4* (ACD Biotechne, 534011) and *Areg* (ACD Biotechne, 430501-C3) were applied to the tissues and incubated at 40°C for 2 hours. Amplification reagents were individually incubated (Amp1, 30 minutes; Amp2, 30 minutes; Amp3, 15 minutes) and washed with wash buffer. Probes were detected individually using HRP-C1 (for *Muc4*) and HRP-C3 (for *Areg*) detection reagents, each incubated for 15 minutes. After washing, Vivid fluors were diluted 1:750 in TSA buffer reagent and applied for 30 minutes. After washing, HRP blocker was applied for 15 minutes to complete signal development. Tissues underwent final washing and DAPI (Fisher, EN63348) was applied for 30 seconds to stain nuclei. After washing in water, tissues were mounted with ProLong Diamond Antifade Mountant (Fisher, P36961).

Pathology scoring

Hematoxylin and eosin-stained slides were observed by a board-certified veterinary pathologist (VVP) using an Olympus BX46 microscope and images were acquired with an Excelis 4K camera. Scoring was performed in a blinded fashion. In C57BL/6 mice, initial observations did not reveal any substantial pathology features such as hyperplasia, metaplasia, or dysplasia. Therefore, those tissues were scored for two parameters as follows. Inflammation: 0 indicates no inflammation, 1 indicates scattered mononuclear cells with or without polymorphonuclear cells in the mucosa, 2 indicates aggregates of inflammatory cells in the mucosa and submucosa, 3 indicates organizing nodules of lymphocytes in the submucosa/mucosa, and 4 indicates inflammatory cells extending through the muscularis. Limiting ridge inflammation: 0 indicates no inflammation, 1 indicates scattered mononuclear/polymorphonuclear cells in the submucosa adjacent to the limiting ridge but glands unaffected, 2 indicates aggregates of mononuclear/polymorphonuclear cells with glands minimally affected, and 3 indicates aggregates of mononuclear/polymorphonuclear cells with glands either affected or lost. For *Mist1-Kras* mice, a previously established scoring system [47] was used with the following modifications. Inflammation: 1 indicates multifocal aggregates of mononuclear ± polymorphonuclear leukocytes in the mucosa, 2 indicates aggregates of inflammatory cells in the submucosa ± mucosa, 3 indicates organizing nodules of lymphocytes and other inflammatory cells in the submucosa ± mucosa, 4 indicates sheets of inflammatory cells extending into or through the muscularis ± adventitia. Pseudopyloric metaplasia: 1 indicates replacement of <25% of oxyntic mucosa by surface-type or antralized gastric glands, 2 indicates replacement of 25-50% of oxyntic mucosa by surface-type or antralized gastric glands, 3 indicates replacement of 50-75% of oxyntic mucosa by antralized glands, 4 indicates >75% replacement of zymogenic glands by antralized mucosa. The individual histology sub-scores are presented in Figure S4 and the proportion of mice exhibiting each score is presented in Figure 4.

DNA and RNA isolation

RNA was extracted from two types of stomach tissue. After the stomach was harvested, the forestomach region was excluded; the remaining glandular stomach was flattened and longitudinally sectioned into three equal parts, each containing corpus and antrum. One-third of this material was frozen on dry ice and stored at -80°C, then used for RNA extraction. As well, a 2-millimeter biopsy was collected from the stomach corpus and stored in 1 ml TRIzol (Thermo Fisher Scientific; 15596026) at -80°C. For RNA extraction from the one-third portion, samples were transferred to an M tube (Miltenyi, 130-096-335) in 1 ml TRIzol (Thermo Fisher Scientific; 15596026) and homogenized using the "M Tube RNA_01.01" program (run for 55 seconds, then stop) in a gentleMACS Dissociator (Miltenyi, 130-093-235). RNA was extracted from the homogenate using chloroform and isopropanol in accordance with the TRIzol Reagent User Guide. DNA was extracted from the aqueous phase in accordance with the TRIzol Reagent User Guide. For RNA extraction from the biopsy punch, samples were ground in TRIzol with a pestle (SP Bel-Art; F65000-0006), then the RNA was extracted with chloroform and precipitated with isopropyl alcohol. RNA pellets were washed with 70% ethanol and then resuspended in RNase-free water.

qRT-PCR for gene expression analysis

Eight mice per treatment group were randomly selected for gene expression analysis. cDNA was constructed with the HighCapacity cDNA Reverse Transcription kit (ThermoFisher Scientific, 4368814) according to the manufacturer's protocol. Gene expression was assessed using RT-qPCR with SYBR Green (Applied Biosystems, A46112) on a QuantStudio 6 Real-Time PCR system (Applied Biosystems, 4485691). Gene expression was normalized to the average of *Hprt*, *Ppia*, *H2afz*, *β-actin*, and *Tbp* housekeeping genes using the ΔΔCt method. Primer sequences are listed below. Some data points were considered biological outliers; however, RNA integrity, amplification efficiency, and melting curve analysis confirmed technical validity, so these data points were not excluded from analysis.

qPCR for *Hp* quantification

Six mice per treatment group were used for *Hp* quantification using DNA extracted from stomach-thirds via the TRIzol method (see above). Copies of the *Hp ureC* gene were measured using qPCR with SYBR Green (Applied Biosystems, A46112) on a QuantStudio 6 Real-Time PCR system (Applied Biosystems, 4485691). Copies were normalized to the average copies of *Ppia*, *H2afz*, and *Tbp* housekeeping genes using the ΔCt method. For data visualization, *ureC* copies were shown relative to $10^6$ mouse housekeeping copies. Primer sequences are listed below.

Primer sequences

| Primer | Sequence (5' → 3') |
|---|---|
| Cldn7-F | GGCCTGATAGCGAGCACTG |
| Cldn7-R | GTGACGCACTCCATCCAGA |
| Reg3g-F | ATGCTTCCCCGTATAACCATCA |
| Reg3g-R | GGCCATATCTGCATCATACCAG |
| ureC (glmM)-F | AAGCTTACTTTCTAACACTAACGC |
| ureC (glmM)-R | GTTTGTGAGCGAATGCATGC |
| Muc2-F | ATGCCCACCTCCTCAAAGAC |
| Muc2-R | GTAGTTTCCGTTGGAACAGTGAA |
| Muc6-F | ACATTGAAGCAAGAGTCTAAGGG |
| Muc6-R | GTGGTACTGTTGGTCTCCGT |
| Muc13-F | GATCTCTGCAACCCTAACCCC |

| | |
|---|---|
| Muc13-R | TCCTTTCACACATGACGACAG |
| Muc4-F | CCTCCTCTTGCTACCTGATGC |
| Muc4-R | GGAACTTGGAGTATCCCTTGTTG |
| Cd44v10-F | CCCAGTGACCCCTGCTAAAA |
| Cd44v10-R | CGCACTTGAGTGTCCAGCTA |
| Reg1-F | ATGGCTAGGAACGCCTACTTC |
| Reg1-R | CCCAAGTTAAACGGTCTTCAGT |
| Mmp10-F | GAGCCACTAGCCATCCTGG |
| Mmp10-R | CTGAGCAAGATCCATGCTTGG |
| Hprt-F | TCAGTCAACGGGGGACATAAA |
| Hprt-R | GGGGCTGTACTGCTTAACCAG |
| Ppia-F | GAGCTGTTTGCAGACAAAGTTC |
| Ppia-R | CCCTGGCACATGAATCCTGG |
| H2afz-F | CCAAGACAAAGGCGGTTTCC |
| H2afz-R | TCCTGCCAACTCAAGTACCTC |
| βtubulin-F | GATCGGTGCTAAGTTCTGGGA |
| βtubulin-R | AGGGACATACTTGCCACCTGT |
| Tbp-F | AGAACAATCCAGACTAGCAGCA |
| Tbp-R | GGGAACTTCACATCACAGCTC |

RNA-sequencing

Four C57BL/6 mice per treatment group (+/- high-fat diet; +/- *H. pylori* infection) were used. Total RNA was extracted from one-third portions of the stomach (corpus and antrum) as described above and RNA was processed for RNA-seq by Novogene (Sacramento, CA, USA). Magnetic beads conjugated with poly-T oligonucleotides were used to purify messenger RNA (mRNA) from total RNA. Library construction was completed using the VAHTS Universal V10 RNA-seq Library Prep Kit for Illumina. The libraries were subsequently quantified using Qubit and real-time quantitative PCR (qPCR), and fragment size distribution was assessed using a Bioanalyzer. Upon passing quality control, the libraries were pooled based on effective concentration and target data output, followed by Illumina sequencing on a NovaSeq X Plus using the Sequencing by Synthesis method.

Analysis was conducted by the Collaborative Core for Cancer Bioinformatics in the Purdue Institute for Cancer Research. All 16 samples were included in the initial analysis. The program fastp v0.23.2 [52] was used to trim reads based on sequence quality and to remove adapter sequences. STAR v2.7.11.b [53] was used to map data to the *Mus musculus* GRCm39 reference genome. FeatureCounts within the Subread package v2.0.1 [54] was used to count reads mapping to genes. The Bioconductor package edgeR v4.0.16 [55, 56] was used within R v4.3.3 to perform statistical analyses, including all differential expression analyses. Because principal component analysis showed that one mock-infected, high-fat diet sample clustered with the standard diet samples and vice versa (as seen in Figure 2A), these two samples were excluded from subsequent analyses. Differential gene expression was determined using a cutoff significance level of false discovery rate ($p_{adjusted}$) < 0.05. RStudio software was used for data visualization. Pathway analysis was performed using Ingenuity Pathway Analysis (IPA, Qiagen). For the diet—interaction analysis, differential expression was assessed using generalized linear models implemented in edgeR. A diet × infection interaction term was tested within stomach samples using quasi-likelihood F-tests, with mock samples serving as the uninfected reference condition. For stomach tissue, the following was computed:

$$logFC = \frac{(\text{Stomach\_HighFatDiet\_Infected} - \text{Stomach\_ HighFatDiet \_Mock})}{(\text{Stomach\_StandardDiet\_Infected} - \text{Stomach\_StandardDiet\_Mock})}$$

Results were interpreted as follows: Positive indicates stronger infection response under high-fat diet; negative indicates stronger infection response under standard diet; ~0 indicates additive effects (infection response is additive and independent of diet).

Statistical analysis

RNA-seq data were analyzed as described above. For qRT-PCR and pathology scoring, statistical analyses were conducted in GraphPad Prism v10.5.0. No mice were excluded from qRT-PCR analyses; one mouse was excluded from pathology scoring analyses as the tissue section was incomplete and could not be fully evaluated. Data were analyzed by non-parametric statistical methods, *i.e.*, the Mann-Whitney U test. For data visualization, bars show the median values and error bars show the interquartile range, as nonparametric summary statistics were used due to the presence of biological outliers. $P < 0.05$ was considered statistically significant.

Data availability

RNA-seq data have been deposited at GEO and will be released from embargo at the time of publication. All data, metadata, and methods used to reach the conclusions in the paper and any additional data required to replicate the study findings will be provided upon request.

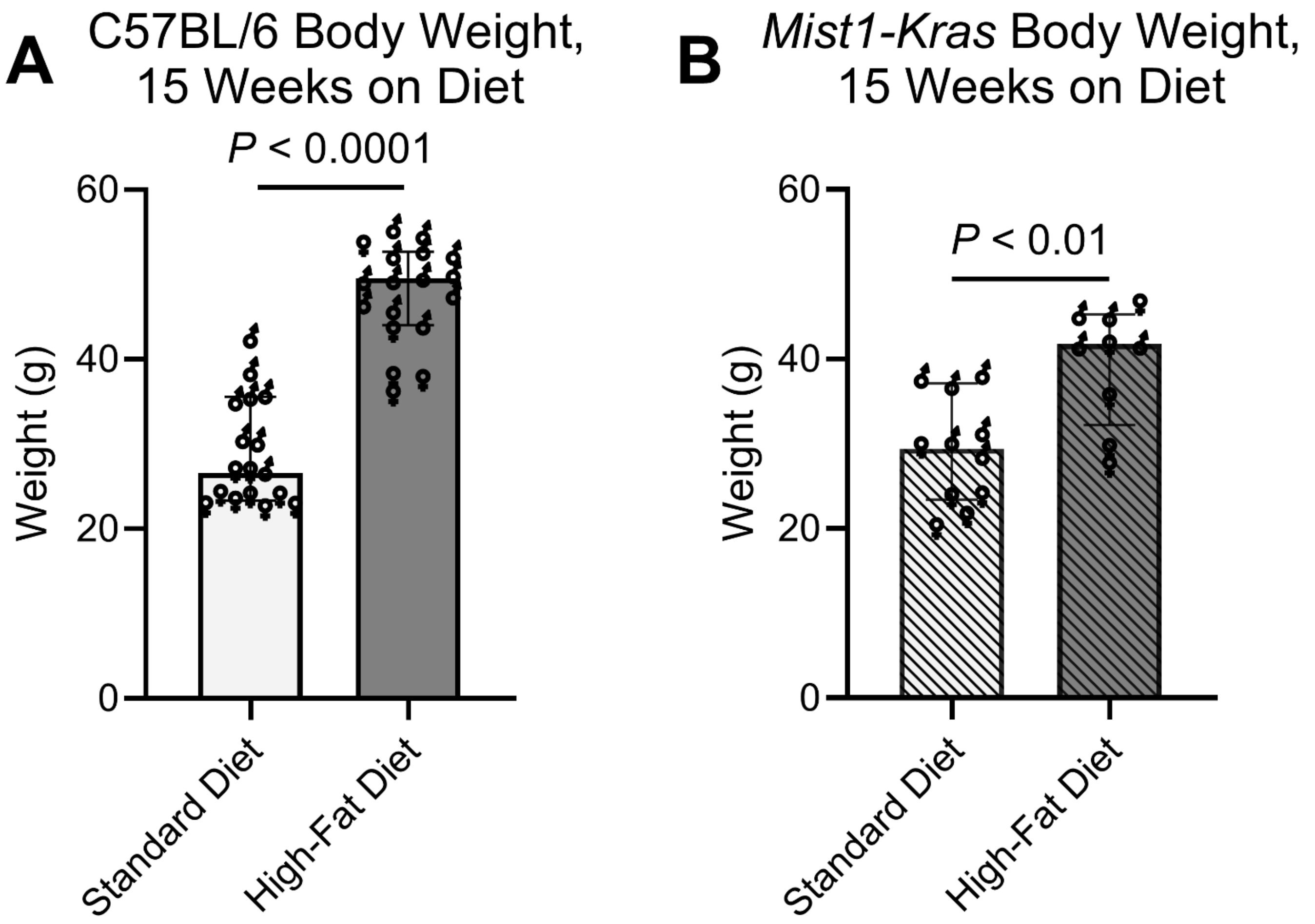


**Figure S1. High-fat diet rapidly induces weight gain in C57BL/6 and *Mist1-Kras* mice.** Mice were randomly assigned to receive a standard diet (10% kilocalories from fat) or high-fat diet (60% kilocalories from fat) for 15 weeks. Body weights at this time point are shown for C57BL/6 mice (**A**) and *Mist1-Kras* mice (**B**). At this stage of the experiment, mice were randomly assigned to be infected with *Hp* or mock-infected. Statistical significance was assessed using the Mann-Whitney U test. Dots represent individual values for each mouse (♂, male; ♀, female), bars indicate the median and error bars indicate the interquartile range. Data for each graph are from N=1 mouse experiment with n=17-18 mice per group (**A**) or n=9-11 mice per group (**B**).

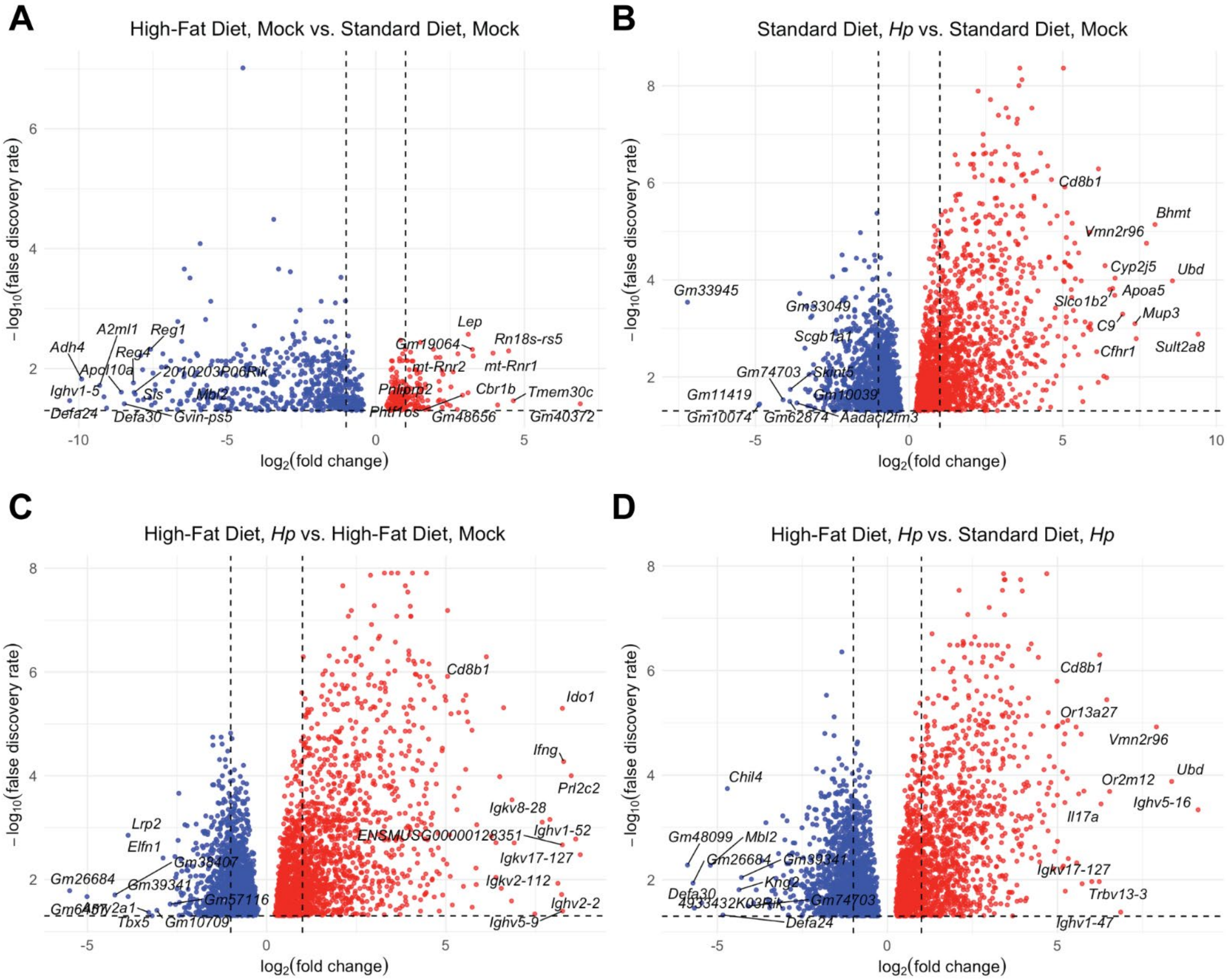


**Figure S2. *Hp* had a greater impact on gastric gene expression than diet did.** Shown are volcano plots for the indicated comparisons. The x-axis indicates the magnitude of the differential gene expression and the y-axis indicates the magnitude of the statistical significance. Red (right side of plots) indicates upregulated genes and blue (left side of plots) indicates downregulated genes. **A)** The top 10 genes by fold change are labeled, along with *Lep*, *Pnliprp2*, *Defa24*, *Defa30*, *Mbl2*, *Reg1* and *Reg4*. **B)** The top 10 genes by fold change are labeled, along with *Cd8b1*, *C9* and *Cfhr1*. **C)** The top 10 genes by fold change are labeled, along with *Cd8b1* and *Ifng*. **D)** The top 10 genes by fold change are labeled.

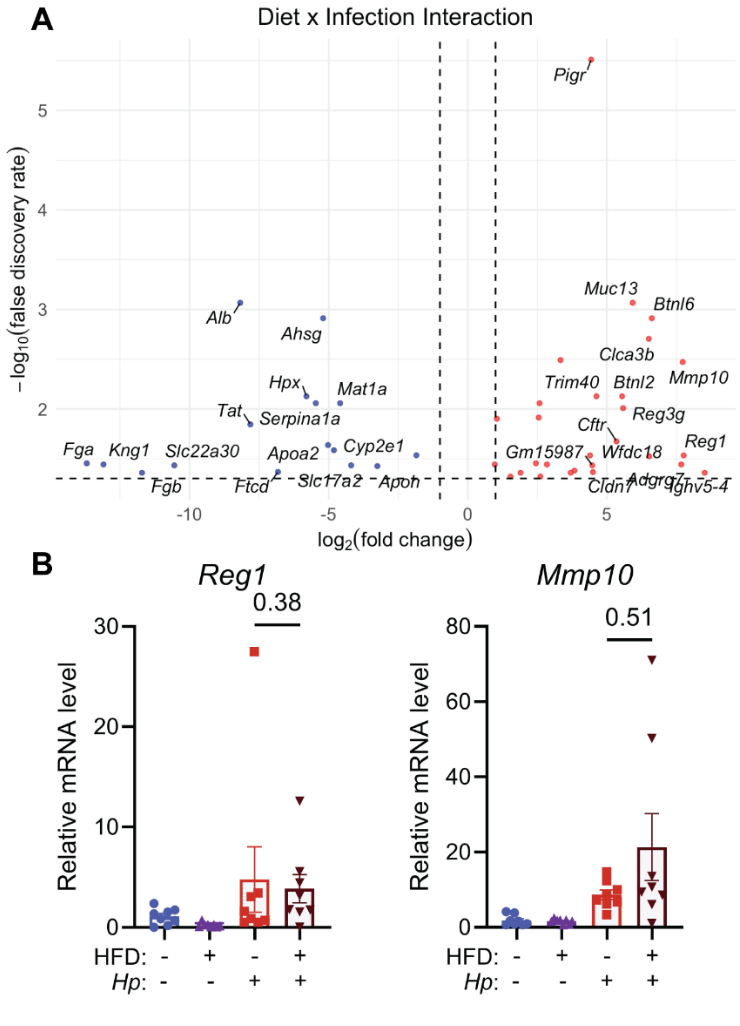


**Figure S3. A unique gene expression signature reflects *Hp* infection during high-fat diet exposure. A)** In the volcano plot, the x-axis indicates the magnitude of the differential gene expression and the y-axis indicates the magnitude of the statistical significance. Red (right side of plot) indicates genes with a stronger infection response in mice fed a high-fat diet and blue (left side of plot) indicates genes with a stronger infection response in mice fed a standard diet. The top 15 genes by fold change are labeled. **B)** The expression of the indicated genes was assessed in additional mice by qRT-PCR using RNA extracted from corpus biopsy punches. Statistical significance was assessed using the Mann-Whitney U test. Dots represent individual values for each mouse, bars indicate the median and error bars indicate the interquartile range. Data are from n=13-18 mice per group assessed in N=2 independent mouse experiments; n=8 mice per group were chosen randomly for qRT-PCR.

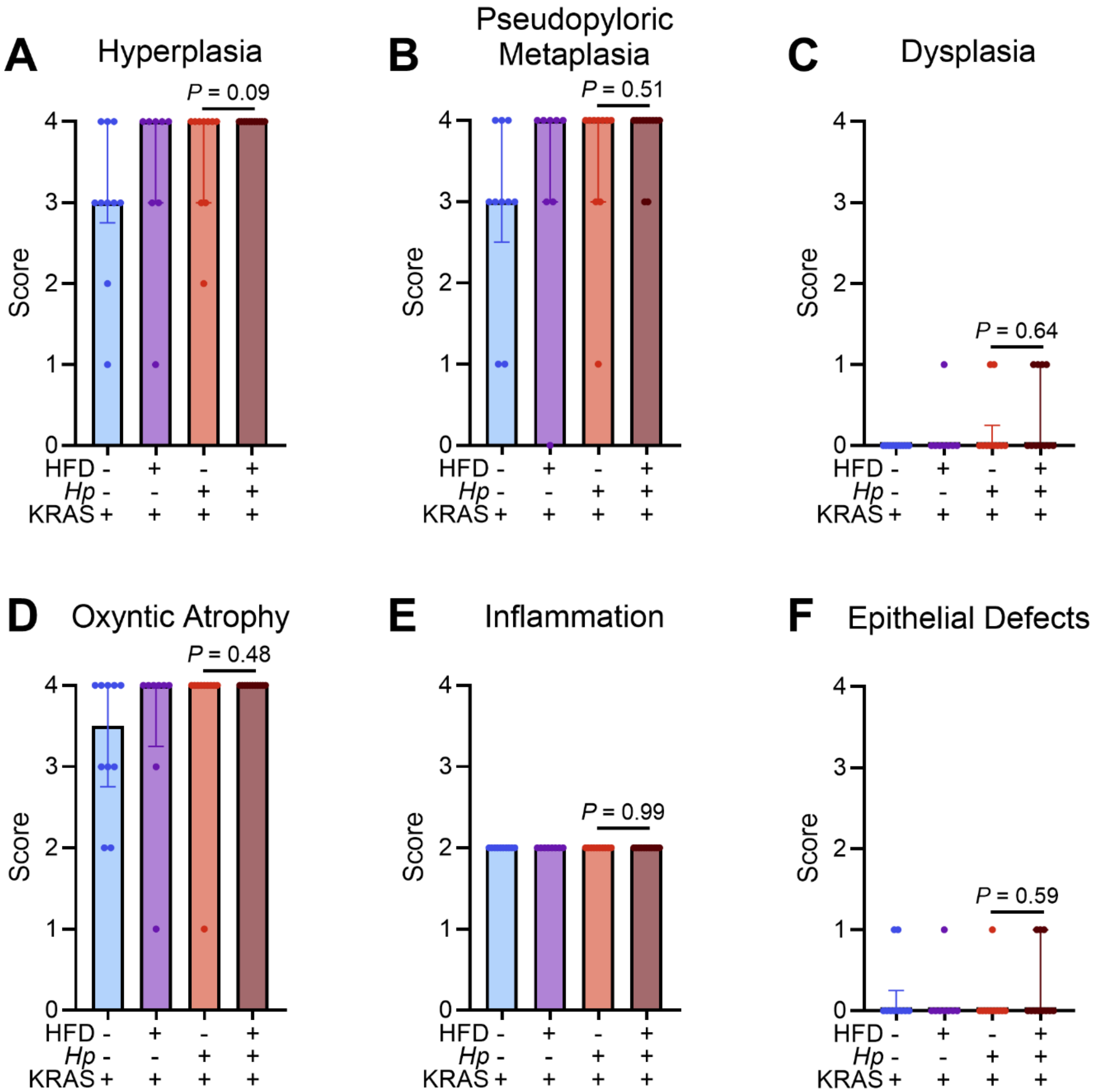


**Fig S4. High-fat diet increases hyperplasia in *Hp*+KRAS+ mice.** A blinded analysis of tissue pathology was performed by a board-certified veterinary pathologist using the modified Rogers criteria (see Methods). **A-F)** The individual parameters that are evaluated in the Total Histology Score are shown. Statistical significance was assessed using the Mann-Whitney U test. Dots represent individual values for each mouse, bars indicate the median and error bars indicate the interquartile range. Data are from n=8-11 mice per group assessed in N=2 independent mouse experiments. **A-D)** Individual datapoints from the contingency tables in Figure 4 are shown, whereas Figure 4 shows the proportion of mice exhibiting each score. **E-F)** Individual datapoints for each mouse are shown; although diet and infection did not impact pathology scores for inflammation or epithelial defects, these metrics are still factored into the Total Histology Score shown in the main text.

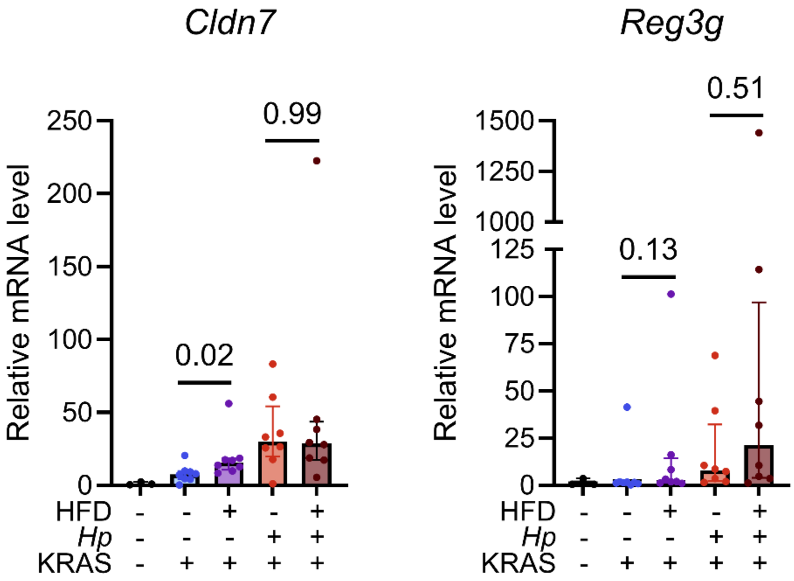


**Figure S5. Metaplasia impacts *Cldn7* expression.** Expression of the indicated genes was assessed using qRT-PCR of RNA extracted from gastric corpus biopsy punches, relative to corpus biopsy RNA from n=3 healthy control mice (white bars, black dots). Statistical significance was assessed using the Mann-Whitney U test. Dots represent individual values for each mouse, bars indicate the median and error bars indicate the interquartile range. Data are from n=8-11 mice per group assessed in N=2 independent mouse experiments; n=8 mice per treatment group were chosen randomly for qRT-PCR. Note that the y-axis for *Reg3g* has two segments to account for an outlier mouse.

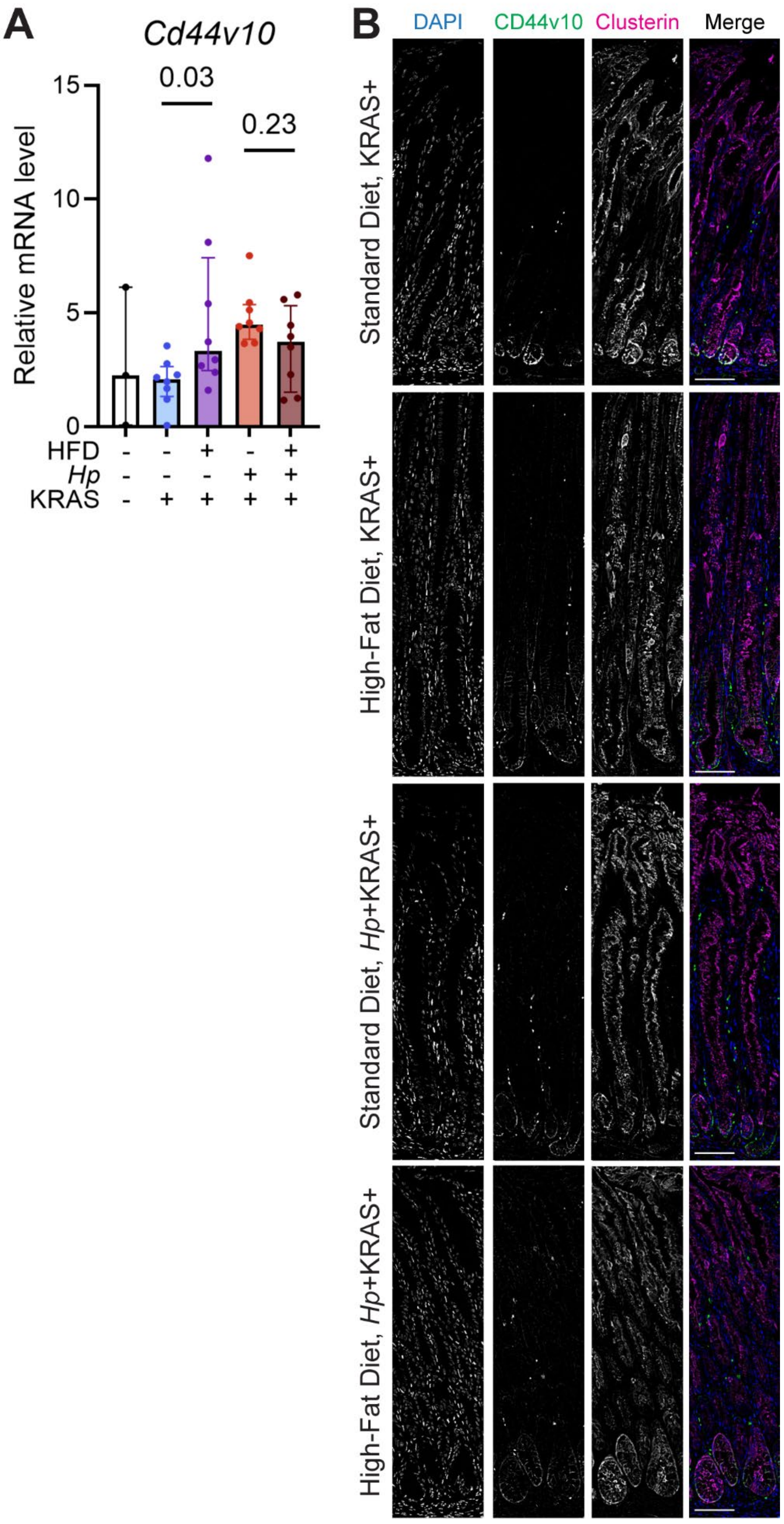


**Figure S6.** High-fat diet impacts expression of the pseudopyloric metaplasia marker CD44v10. **A)** Expression of *Cd44v10*, orthologous to human *CD44v9*, was assessed using qRT-PCR of RNA extracted from gastric corpus biopsy punches, relative to corpus biopsy RNA from n=3 healthy control mice (white bars, black dots).

Statistical significance was assessed using the Mann-Whitney U test. Dots represent individual values for each mouse, bars indicate the median and error bars indicate the interquartile range. Data are from n=8-11 mice per group assessed in N=2 independent mouse experiments; n=8 mice per treatment group were chosen randomly for qRT-PCR. **B)** Immunofluorescence microscopy was used to detect gastric corpus expression of CD44v10 (green in merge) and the metaplasia marker clusterin (magenta in merge). DAPI (blue in merge) was used to stain nuclei. Representative images are shown. Scale bars, 100 µm.
